# The neurodevelopmental genes *alan shepard* and *Neuroglian* contribute to female mate preference in African *Drosophila melanogaster*

**DOI:** 10.1101/2022.05.15.491994

**Authors:** Paula R. Roy, Dean M. Castillo

**Author notes:** Current affiliation: Department of Neurobiology, University of Utah, Salt Lake City UT. Corresponding Author, Current affiliation: Department of Biological Sciences, Miami University, Hamilton OH.

## Abstract

Mate choice is a key trait that determines fitness for most sexually reproducing organisms, with females often being the choosy sex. Female preference often results in strong selection on male traits that can drive rapid divergence of traits and preferences between lineages, leading to reproductive isolation. Despite this fundamental property of female mate choice, very few loci have been identified that contribute to mate choice and reproductive isolation. We used a combination of population genetics, quantitative complementation tests, and behavioral assays to demonstrate that *alan shepard* and *Neuroglian* contribute to female mate choice, and could contribute to partial reproductive isolation between populations of *Drosophila melanogaster*. Our study is among the first to identify genes that contribute to female mate preference in this historically important system, where female preference is an active premating barrier to reproduction. The identification of loci that are primarily known for their roles in neurodevelopment provides intriguing questions of how female mate preference evolves in populations via changes in sensory system and higher learning brain centers.

## Introduction

For sexually reproducing organisms mate choice is critical to balance maximizing fitness with avoiding costly heterospecific mating events (Mendelson and Shaw, 2012; Gray, 2022). Strong selection on male courtship traits and female preferences can drive rapid divergence between closely related lineages (Fisher, 1958; Andersson, 2019), providing a link between sexual selection and the evolution of reproductive isolation. A large body of work has focused on documenting and describing female preference and identifying genes controlling male courtship traits (Chenoweth and Blows, 2006; Yamamoto and Koganezawa, 2013; Neelon et al., 2019; Munson et al., 2020). However, there have been fewer studies that have identified specific genes that contribute to female mate preference, with most instead focusing on quantitative trait loci (Bay et al., 2017; Svensson et al., 2017; Blankers et al., 2019; Xu and Shaw, 2019a; Chowdhury et al., 2020) and some identifying specific genes (Chowdhury et al., 2020). Identifying and characterizing loci contributing to female preference is a critical first step to understand how courtship signal detection, signal processing, and higher brain functions (i.e., learning and memory) contribute to sexual selection and female mate preference evolution.

One challenge in identifying the genetic basis of female choice lies in the complexity of preference-based behaviors (Andersson, 2019). During mate choice females are receiving many, potentially multimodal, signals that then require processing before a decision to accept or reject a mate is made (Gray, 2022; Munson et al., 2020). Progress has been made to understand male courtship and how female preference exerts divergent selection between populations (Xu and Shaw 2019a; Xu and Shaw 2021), yet even in systems where male courtship traits, and presumably male signals, are well understood, identifying female loci can be difficult (Xu and Shaw 2019b). The complexity of female preference implies that this trait is polygenic, but it is unclear how many loci we would expect to contribute to this trait. Specifically, it remains unclear whether multiple loci interact in an emergent fashion, contributing to female preference as a whole, or if individual loci contribute to a specific facet (perception of a specific signal/cue) of female preference (Boake et al., 2002; Chenoweth and Blows, 2006). Additionally, while premating isolation evolves rapidly and is often thought to evolve before other barriers to reproduction (Coyne and Orr, 1989; Turissini et al., 2018), it can be difficult to determine if the behaviors that currently act as a mating barrier contributed to the speciation event or evolved after species divergence (Coyne and Orr, 2004; Safran et al., 2013; Kopp et al., 2018). Focusing on populations that vary in the degree of mating isolation can overcome some of these challenges and can be used as a model to understand the early stages of divergence and identify the loci that contribute to speciation and reproductive isolation.

*Drosophila melanogaster* originated in Southern Africa but is now a cosmopolitan species with a worldwide distribution. This broad range has resulted in structured and cryptic populations, even within Africa (Coughlan et al 2021), and strong premating isolation between some specific populations (Wu et al., 1995). For example, females from strains collected in Southern Africa strongly reject males from strains that are collected from non-African localities in both choice and no-choice mating experiments (Wu et al., 1995; Hollocher et al., 1997; Coughlan et al., 2021). These strong premating isolation behaviors are also observed in other parts of the *D. melanogaster* range including the Southeast United States and the Caribbean (Yukilevich and True, 2008). These mating preferences and behaviors that occur outside of Africa are potentially a product of African ancestry in these populations and admixture with non-African populations (Kao et al 2015; Bergland et al 2016). Combined, these observations of strong and variable female preference and population structure provide an ideal system to explore the evolution of female mate preference.

In this study, we leverage these *D. melanogaster* populations by combining population genomics and behavioral assays to identify genes that contribute to female mate preference and partial reproductive isolation. First, we use previously published genomic data from Kao et al. (2015) to identify loci that show a clinal pattern of allele frequency in an admixed secondary contact zone, suggesting a role in reproductive isolation (Harrison and Larson, 2014). We compare these candidates to outlier loci from other studies allowing us to narrow our list and focus on two loci, *alan shephard* (*shep*) and *Neuroglian* (*Nrg*). We then examine patterns of differentiation in these loci between African and non-African genomes and further provide genetic evidence that these two loci contribute to African female mate preference.

## Methods

### Genomic cline analysis

To understand how alleles potentially involved in reproductive isolation are structured we compared allele frequencies across the Southeastern United States and Caribbean using previously published data (Kao et al 2015). We focused on these populations because this clinal structure allowed us to make explicit expectations for a gene that contributes to reproductive isolation compared to the rest of the genome (Harrison and Larson 2014). Specifically, because previous studies have demonstrated a correlation between latitude and African ancestry, with more African ancestry in southern populations, we predict that genes involved in behavioral isolation will have a steeper slope (i.e., larger regression coefficient) compared to the rest of the genome. Using the genotype matrix (genotype calls for all variable positions they identified) from Kao et al. (2015), we fit linear regression models between SNP allele frequency and latitude for the populations.

#### GO Enrichment Analysis

We retained the top 1% outliers for use in a gene ontology enrichment analysis. The list of genes used for the enrichment analysis are found in Supplemental Table 1. This list included genes that contained outlier SNPs. SNPs that were not within the known boundaries of a gene were not included in this analysis. That is, we did not assign SNPs to the nearest gene. We used the web application FlyEnrichr (Chen et al. 2013; Kuleshov et al. 2016), to look for enrichment in “biological processes”. These biological processes are defined by Flybase (The Gene Ontology Consortium 2017) and FlyEnrichr uses definitions made in the 2018 version of Flybase.

#### Comparison to other datasets

We compared our list of outlier genes to two other datasets that looked for differences in behavior between African and non-African flies using either DNA sequence data or RNAseq transcriptomic data. In the first comparison we compared our set of outliers to Coughlan et al. (2021). In this study the authors compared allele frequency between strains that showed strong female choice and strains that did not show strong choice and identified outlier loci using the PBE test statistic. In the second comparison we compared our list of outliers to Bailey et al. (2011) who looked at transcriptomic data in female brains after they interacted with “preferred” and “non-preferred” males in both African and non-African strains.

### Choice of candidate loci

After calculating regression coefficients for each SNP, we found that two genes previously identified as important for female behavior *alan shepard* (*shep*; Chen et al., 2014) and *Neuroglian* (*Nrg;* Carhan et al., 2005) were genomic outliers (See Results). These genes were outliers in at least one of the additional data sets we compared our lists with (see Results) and appeared as key loci contributing to significant GO terms in the enrichment analysis (See Results). Previous data for these genes demonstrated that null mutant females rejected were mating deficient, and we used these mutants to test for a specific role in African female behavior.

### Population genetics of shep and Nrg across Drosophila populations

Several studies have examined the population genetics of *Drosophila* populations from worldwide distributions (Lack et al. 2016; Kapopoulou et al. 2018; Sprengelmeyer et al. 2020), but our focus was to specifically look at variation in *shep* and *Nrg*. The SNPs identified in Kao et al (2015), the source of our cline analysis, likely represent a subset of the total variation observed in these two loci, since they focused on SNPs that they could define as coming from either the African or European source population. To understand the variation in these alleles and population differentiation we used the PopFly application (Hervas et al. 2017) to provide summary statistics for nucleotide diversity (π) and differentiation (F_st_) for the two populations with the largest number of sequences strains, Raleigh (RAL) and Zimbabwe (ZI) (Lack et al. 2016). We also exported FASTA sequences from PopFly to examine differentiation using principal components analysis. This allowed to visualize and estimate the variation of alleles/haplotypes within and between populations.

### Quantitative complementation test

To determine if *Nrg* and *shep* alleles contribute to the female mate preference of African strains, we used an experimental design similar to a quantitative complementation test combined with a binary mate choice assay (Supplemental Figure 1). Quantitative complementation tests have previously been used for no-choice tests (Chowdhury et al., 2020) and binary choice tests (Comeault et al., 2017). It is important to understand how typical complementation tests are performed to understand how the expected phenotypes, and potential interpretations, used in our experiments differ. In a typical complementation test, the starting point is two parental strains that have extreme divergent phenotypes for a continuous trait and are often the minimum and maximum trait values in a population (reviewed in Mackay 2001; Supplemental Figure 2). The trait of interest may be polygenic and hybrids have an intermediate phenotype. A third strain is used that carries a mutation in the gene of interest or a large chromosomal deletion that contains the gene of interest. Because mutant strains are lab-derived they typically have an average phenotype when compared to the divergent parental strains. In *Drosophila* large deletions and some null mutations are homozygous lethal. As such they must be “balanced” by a chromosome, known as a balancer chromosome, that has several complex inversions that prevents recombination with the homologous chromosome (Muller 1918; Miller et al. 2019). Balancer chromosomes also carry a dominant visible mutation to track the presence/absence of the balancer chromosome. When both wild-type parental strains are crossed independently to the mutant/balancer strain four genotypes are produced and can be phenotyped (Supplemental Figure 1). Two of the genotypes are heterozygous at the gene of interest. They carry either parental allele and the allele from the balancer chromosome. The other two genotypes are hemizygous. They only carry one the parental alleles. The homologous chromosome either lacks the gene of interest or has a non-functional allele. In *Drosophila* balancer chromosomes have a marker with a dominant visible phenotype. This causes the heterozygote individuals, but not hemizygote individuals, to have a phenotype that could affect the trait of interest and must be taken into account during analysis (Turner et al., 2011). The expected phenotypes, and their relationships the the statistical models used to analyse quantitative complementation testes, are described in the Supplemental Information.

The potential phenotypes we would expect to observe in our system differ from the standard quantitative complementation test for one main reason: our mutant/balancer strains have the same phenotype as one set of parental strains (Supplementarty Fig 2). Given the evolutionary history of the lineages we are using, and the genetic resources available, we lack two extreme divergent parental strains with mutant/balancer strains of an intermediate (“average”) phenotype. In our system, strains from Africa are extremely choosy, representing one extreme of the phenotypic distribution (see below). The other extreme, random mating, occurs in all strains found outside of Africa. That means that parental strains that randomly mate and any mutant/balancer strains available all have the same phenotype because they all have non-African alleles. This results in a minimum phenotype (random mating, 50% choosing one male) that is shared with non-African parental strains and the mutant/balancer strain. This changes the expected phenotypes of heterozygous and hemizyogous progeny produced in these crosses as well as the magnitude and nature of the coefficients from binomial regressions compared to a standard quantitiative complementation test (Supplementary Fig 2; Supplemntary Information). Regardless we used the standard experiment design and discuss how this design might be interpreted differently in the context of our system.

**Figure 1.**
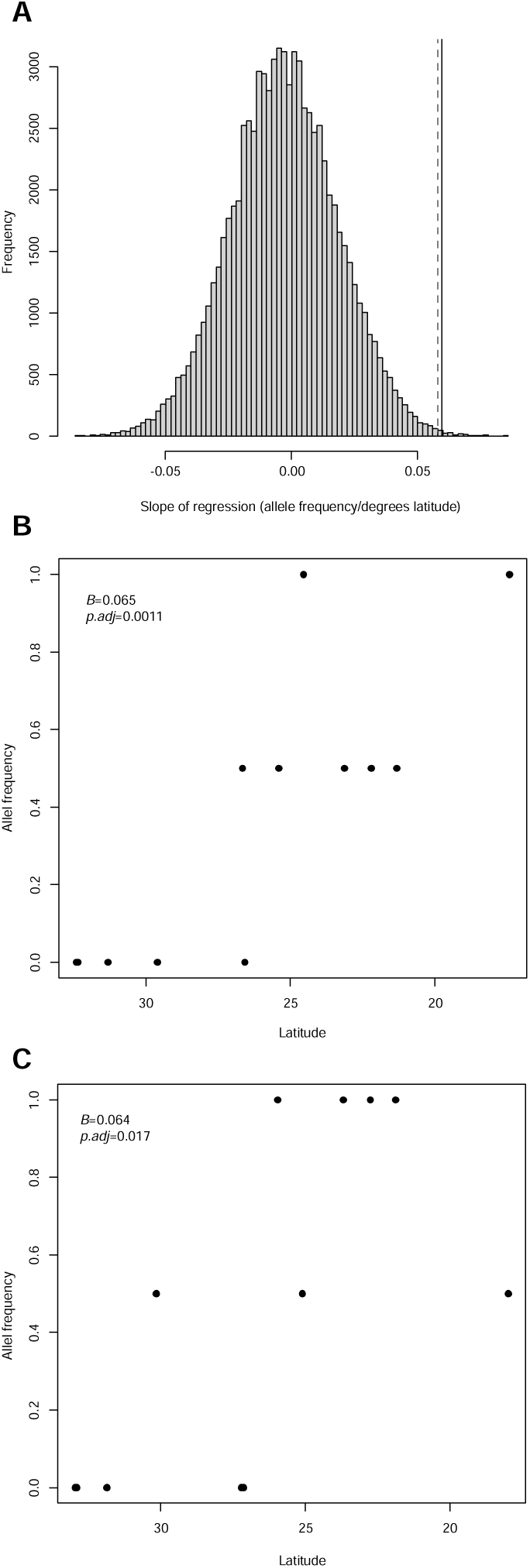
A). The genes *alan shepard* (*shep*) and *Neuroglian* (*Nrg*) are genomic outliers with a stronger relationship in African allele frequency and latitude in an analysis using genomes from populations collected in the Southeastern United States and the Bahamas (data previously collected in Kao et al. (2015)). The dashed line corresponds to the value for *shep* and the solid line corresponds to the value for *Nrg*. The change in allele frequency over latitude for B) *Nrg* and C) *shep*. The regression coefficient and adjusted p-value are provided for each gene.

**Figure 2.**
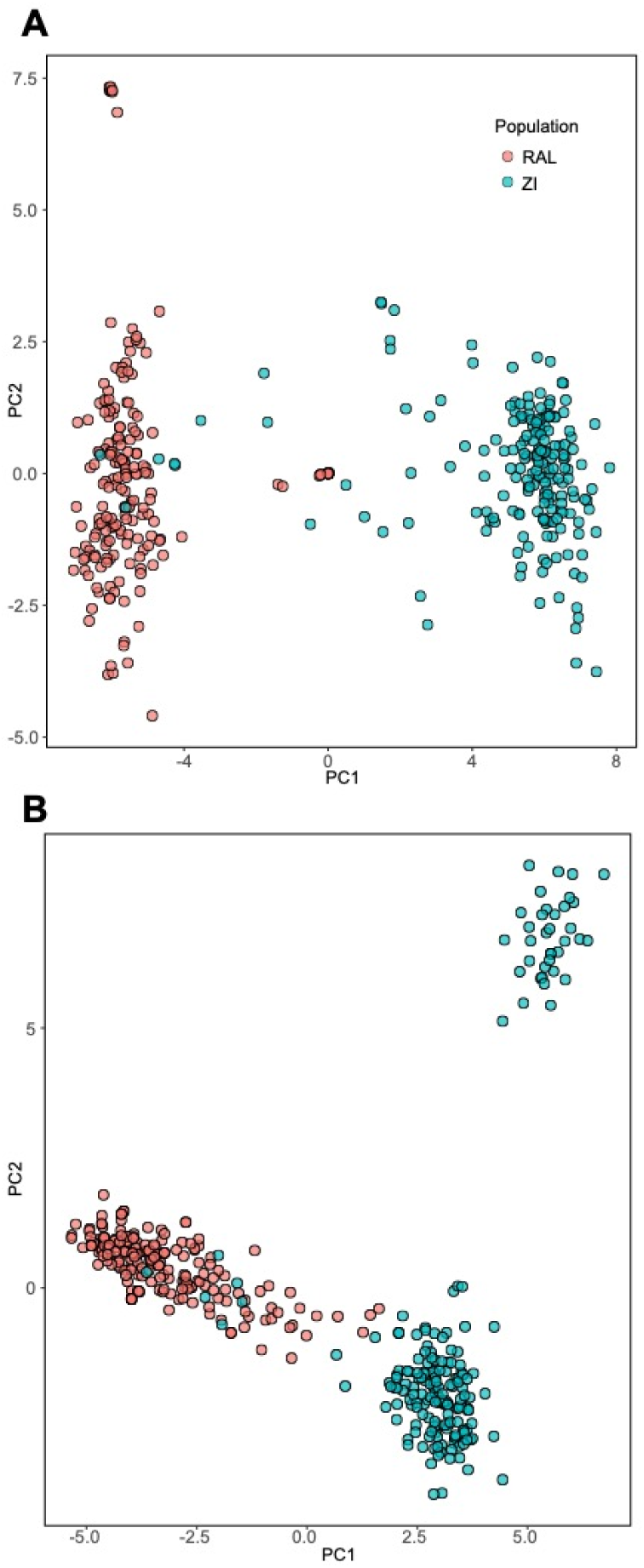
Population differentiation for both A) *shep* and B) *Nrg* are seen in principal components for allelic variation comparing populations from Raleigh NC (RAL, red) and Zimbabwe (ZI, blue).

Using a binary choice test, we introduced a single female of a given genotype (one of the four genotypes produced from the complementation tests crosses) to two males from representative strains: Z53 representing an African male and DGRP882 representing a non-African. Trios were observed for 60 minutes or until copulation occurred. When copulation occurred between a pair of individuals, the non-copulating male was removed and the strain identity was determined (see behavioral assay conditions, below). After the pair was finished copulating, we also examined the successful male to verify which strain the female chose. Our *a priori* expectation was that the females hemizygous for the African allele will prefer African males, whereas females hemizygous for the non-African allele will mate indiscriminately. It has been previously shown that not-mating with either male is a phenotype associated with African female preference (Jin et al 2022). However, in this experiment we noticed this occurred only 3 times total out of all of our replicates with the heterozygote and hemizygote flies.

### Drosophila strains and crosses

To determine whether *Nrg* and *shep* alleles contribute to female preference behavior between populations of *D. melanogaster* we needed to identify wild-type strains that show strong female mate preference and complementary strains that show no preference. Strains from Southern Africa were once considered to be a single lineage from the ancestral range and were referred to as Z-type. All other strains that migrated from the ancestral range (non-African and some Northern African strains) were considered a single cosmopolitan lineage and referred to as M-type. The M-type behavior contrasted with Z-type behavior in that M-type females often mate indiscriminately, showing no preference, whereas Z-type females strongly reject M-type males (Wu et al., 1995). Since this initial description, the demographic and evolutionary relationships between *D. melanogaster* populations have been shown to be more complex (Coughlan et al., 2021), and it is unlikely that all Z-type strains are a homogenous lineage.

We were able to choose strains based on geographic location and evaluated the female preference behavior for wild type females prior to using these strains in our crosses. For our representative African strain, we chose Z53 because its behavior profile is well documented and has been consistent across studies (Wu et al., 1995; Hollocher et al., 1997; Moran, 2006; Jin et al., 2022). For our representative non-African strain, we chose DGRP882 which we had previously used and had expected baseline behavior (Jin et al., 2022). The African admixture has been estimated for individuals of the DGRP panel (Pool, 2015) and DGRP882 has among the least amount of African ancestry making it most likely to be differentiated from African strains. In addition to these representative lines, we also chose two additional African strains: Z30 and a more recently collected strain CH11. Canton-S was used as an additional non-African strain because it is commonly used in behavioral studies (Tompkins et al., 1982; Lasbleiz et al., 2006; Ng and Kopp, 2008; Kohlmeier et al., 2021).

To create heterozygote and hemizygote genotypes for our complementation tests, we chose to use loss of function mutant lines for both *shep* and *Nrg* that had been previously characterized as null alleles (*shep* Chen et al., 2014; *Nrg* Bieber et al., 1989; Enneking et al., 2013). Wild type females that have not mated (called virgins) readily mate with males. *shep* and *Nrg* mutant females that are virgin do not mate and instead show rejection behaviors when courted by males (*shep* Chen et al., 2014; *Nrg* Kerr et al. 1997; Carhan et al. 2005). We used the *shep^BG00836^*null mutation (BDSC #12513), hereafter *shep^-^*. Homozygous null mutant females for this allele are viable and mating deficient. For our complementation test we wanted to use a strain that carried *shep^-^* on one chromosome while the homologous chromosome was a balancer chromosome. We combined the *shep^-^* allele with the TM6B balancer chromosome. This balancer chromosome carries the dominant visible Humoral (*Hu)* mutation. We inbred progeny from this original cross for three generations before using this strain in our behavioral assays. The mutant line and the line that contributed the TM6B chromosome were not the same genetic background so residual heterozygosity could be present in this strain. Any effects of this heterozygosity on the X chromosome and 2^nd^ chromosome would occur in all of the crosses/treatments that we examined.

We used the *Nrg^17^* allele, which is synonymous with *Nrg*^2^ and hereafter *Nrg^-^* (BDSC #5595). This specific *Nrg^-^* allele is homozygous lethal, so the stock is maintained with one X chromosome carrying *Nrg^-^* and the other X chromosome is the balancer FM7 chromosome. Fm7 carries the dominant visible marker *Bar*.

Importantly both the *shep^-^* and *Nrg^-^* strains are non-African and carry non-African alleles of *shep* or *Nrg* on the respective balancer chromosomes. These alleles are not identical to DGRP882 or Canton S alleles, however, all non-African alleles are phenotypically more similar than they are to are to the African alleles, because they result in random mating.

### Experimental Crosses

We crossed the *shep*/TM6B line to both DGRP882 and Z53 males to generate four F1 females. Non-African heterozygotes (N/N) have two non-African *shep* alleles, one from the TM6B chromosome and one from the DGRP882 chromosome. Non-African hemizygotes (N/N-*shep^-^*) only have the non-African *shep* allele. We designate the *shep^-^* chromosome as N-*shep^-^* to highlight that the rest of the chromosome carries non-African (N) alleles. We expect both the N/N and N/N-*shep^-^*strains to have random mating phenotypes. This is a notable difference from a standard quantitative complementation test where these genotypes would show phenotypic differences (Mackay et al 2001; Supplemental Fig 2). African heterozygotes (A/N) have an African allele from the Z53 chromosome and a non-African allele from the TM6B chromosome. African hemizygotes (A/N-*shep^-^*) only carry the African allele of *shep*. If *shep* contributes to female mate preference we would expect to see a significant difference in the A/N-*shep^-^* genotype both in terms of its difference from random mating and from the A/N genotype.

We crossed the *Nrg^-^*/FM7 line to DGRP882 and Z53 similar to the design for *shep.* For experiments with *Nrg,* we tested two additional African and one non-African strains (described above) for two reasons. First, we were not sure how or if the *Bar* locus, which affects eye morphology, would affect female mating behavior. This is particularly important because the impact of vision for female mating behavior could vary between strains. We tested the effect of *Bar* explicitly by including Canton S as an additional strain that should not exhibit strong mating preference. Second, even though our cline analysis identified *Nrg* as a strong outlier, this locus was only a marginal outlier in and independent study that identified putative female preference loci (Coughlan *et al*., 2021). This could indicate that the effect of *Nrg* differs between strains and populations, or multiple alleles of *Nrg* segregate in African populations with only a subset contributing to female preference phenotypes. Either scenario could result in a situation where there is not a single high frequency allele contributing to female preference in African strains motivating us to test multiple African strains for this locus. Our expected results for *Nrg* were the same as for *shep*. Specifically, we expect both the N/N and N/N-*Nrg^-^* strains to have random mating phenotypes. If *Nrg* contributes to female mate preference we would expect to see a significant difference in the A/N-*Nrg^-^* genotype both in terms of its difference from random mating and from the A/N genotype.

### Behavioral assay conditions

All stocks and virgins were kept on standard cornmeal molasses media in an incubator on a 12:12 light cycle held at a constant 20 degrees Celsius and 50% humidity. Virgin males and females were collected within the first 0-4 hours post eclosion to ensure they had not mated. Females were phenotyped based on the presence of the dominant visible marker and housed in groups of 5-10. Males were housed individually because group housing can potentially change courtship vigor (Dixon *et al*. 2003). All individuals were allowed to age 7-10 days before experiments because this is the timeframe that produces the maximum number of matings for the African strains (Jin et al 2022). All behavioral assays were performed in a room held at a constant 20C degrees Celsius and 50% humidity within two hours of the incubator lights turning on to maximize the number of copulations we could observe.

The strains used in the experiment are phenotypically indistinguishable, so to be able to tell males apart we placed them on food containing blue food coloring 48 hours prior to the behavioral assay. This is a non-invasive, robust identification method (Wu et al., 1995; Hollocher et al., 1997). In these studies, all males received food dye treatment, but in our experiment, we only marked one male per mating with blue food dye. We found that we could more easily distinguish between a dyed and non-dyed male rather than distinguishing between males fed two different colors. We tested the effect of this treatment on female preference and found no effect (Supplementary Information).

We used males from the representative strains Z53 and DGRP882 for all binary choice tests. We collected all genotypes for a given locus in each block of our trials in equal numbers, but due to differences in the available individuals after aging, our final number of females was different for each genotype. While the rearing conditions and behavior rooms conditions were very well controlled, our behavioral protocol also included completing a set of DGRP882 females in parallel to our experimental crosses. For each block of trials, 20 DGRP882 females were used as a behavioral control and we consistently had 8-12 (representing 40-60%) choosing Z males consistent with the random mating for this genotype, allowing us to conclude that the environment was not affecting our matings and pool replicates over blocks.

### Statistical analysis of behavioral data

To analyze data from a quantitative complementation test using continuous traits, a linear model of the following structure is used (Equation 1; Pasyukova and Mackay 2000; Mackay 2001). The goal is to isolate the effects of the gene of interest on the phenotype while taking into account the difference in parental strains and any effects the balancer chromosome might have on the phenotype.

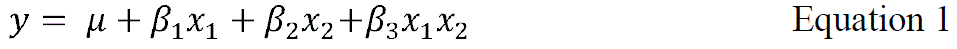

A full description of this analysis framework is included in the Supplemental Information.

For our behavioral preference experiments the data can be represented as binary choices. An individual female replicate chooses either the DGRP882 or the Z53 male, which can be represented as 0 and 1. Our statistical philosophy is to construct a model based on the properties of the data (Warton et al. 2016) and would lead us to using binomial regression. One benefit of the generalized linear models is that the same model structure can be used as Equation 1, except now the scale is in log-odds due to the logit transformation of the data. The logit link function is the most common link function for binomial regression (Equation 2).

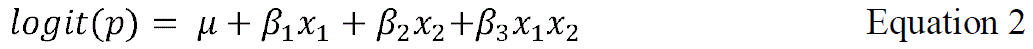

While the model in Equation 2 can be used to describe our data, there are some important differences to consider. Specifically, because the mutant/balancer strain and the non-African parental strain all carry non-African alleles, the genotypes between this strain will have random mating phenotypes leading to the, *β*_1_ coefficient being equal to 0. This coefficient represents the hemizygote effect and is estimated using the N/N-shep-strain (or N/N-Nrg-) data. The *β*_3_ coefficient can still be interpreted as an interaction effect that represents the change in phenotype for the hemizygote across different strains, but this is not the only interpretation or way to statistically model this difference. The, *β*_3_ coefficient can be estimated and interpreted as the difference of the African hemizygote (A/N-shep-) from the African heterozygote (A/N).

Since we do not expect an overall hemizygote effect, and may not see a strain effect, we can propose a model that does not contain a statistical interaction. One strategy to test the inclusion of main vs interaction effects could be sequential likelihood ratio tests. However, because all of our variables are categorial we cannot drop a main effect from a model while retaining an interaction term. Doing this would result in a reparameterization of the model that still contain four coefficients. In light of this observation, we can reparametrize our model purposefully so that regression coefficients represent the genotype differences that we care about. In this model, instead of the African hemizygote being represented by the interaction term and being coded as x1×2 (Equation 2) we can capture the effects of that genotype as its own variable x3 (Equation 3).

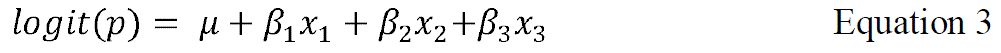

Comparing the model in Equation 2 to the model in Equation 3 we estimate the same number of coefficients. We can now model the trait mean of each genotype independently, and do not expect any covariance in our coefficient estimates, each coefficient is independent. If a candidate gene is important for female preference we would expect to see a significant difference in the African hemizygote (A/N-*Nrg^-^*) genotype both in terms of its difference from random mating and from the A/N genotype. To test this contrast explicitly we used a Wald’s test to test for significant difference between β_2_ and β_3_ coefficients in Equation 3. This would indicate a significant shift in preference for the African hemizygote genotype equivalent to what we would observe if we combined our coefficients from the interaction model to estimate the effect of this genotype (Supplementary Information). This approach is equivalent to several studies that compare heterozygotes and hemizygotes generated from a single parent strain directly to each other to asses gene function either using large sets of deficiencies (Takahashi et al. 2011; Lee et al. 2016) or candidate genes (Balakireva et al. 1998; Matsuo et al. 2007; Comeault et al 2017). This last study, Comeault et al. (2017), has a similar design to ours where the mutant/balancer strain has the same phenotype of one of the parental strains. They use a binomial regression that directly compared heterozygotes and hemizygotes equivalent to the model we propose in Equation 3 (Supplemental Information).

In the results we report both the interaction models and the models from Equation 3. For interaction models we report coefficients and their corresponding p-values to remain consistent with other complementation test studies. For this second model (Equation 3) we calculated 95% confidence intervals and determined whether any interval contained 0. We also provide results for the Wald’s test specifically testing for differences in African heterozygote and hemizygote genotypes.

## Results

### Female behavior loci are genomic outliers

To generate candidate genes for functional validation, we leveraged genomic data from populations of *D. melanogaster* that exhibit a cline in African ancestry (Kao et al., 2015; Bergland et al., 2016) and African-like female preference behavior (Yukilevich and True, 2008). Our goal was to identify outlier genes that showed a steeper cline in African ancestry compared to the genome wide average. We used these genes to understand patterns of gene ontology enrichment and compared our list of candidates to independent analyses that used populations from Africa to identify female preference alleles (Coughlan et al., 2021) and changes in gene expression in female brains between these populations (Bailey et al. 2011).

We retained the top 1% of outliers for loci that had the steepest clines with African alleles being more frequent at lower latitudes. We determined whether SNPs occurred in annotated genes, and retained these genes for a gene ontology (GO) enrichment analysis (Supplemental Table 1). We focused on biological function and observed a pattern where axon and neuron maintenance and development were enriched in our top 10 categories (Table 1). Courtship behavior also showed significant enrichment, and within that list were two genes, *alan shepard* (*shep*) and *Neuroglian* (*Nrg*) that contribute specifically to female behavior. Interestingly *Nrg* shows up as a gene contributing to many of the GO terms that show significant enrichment.

**Table 1.**
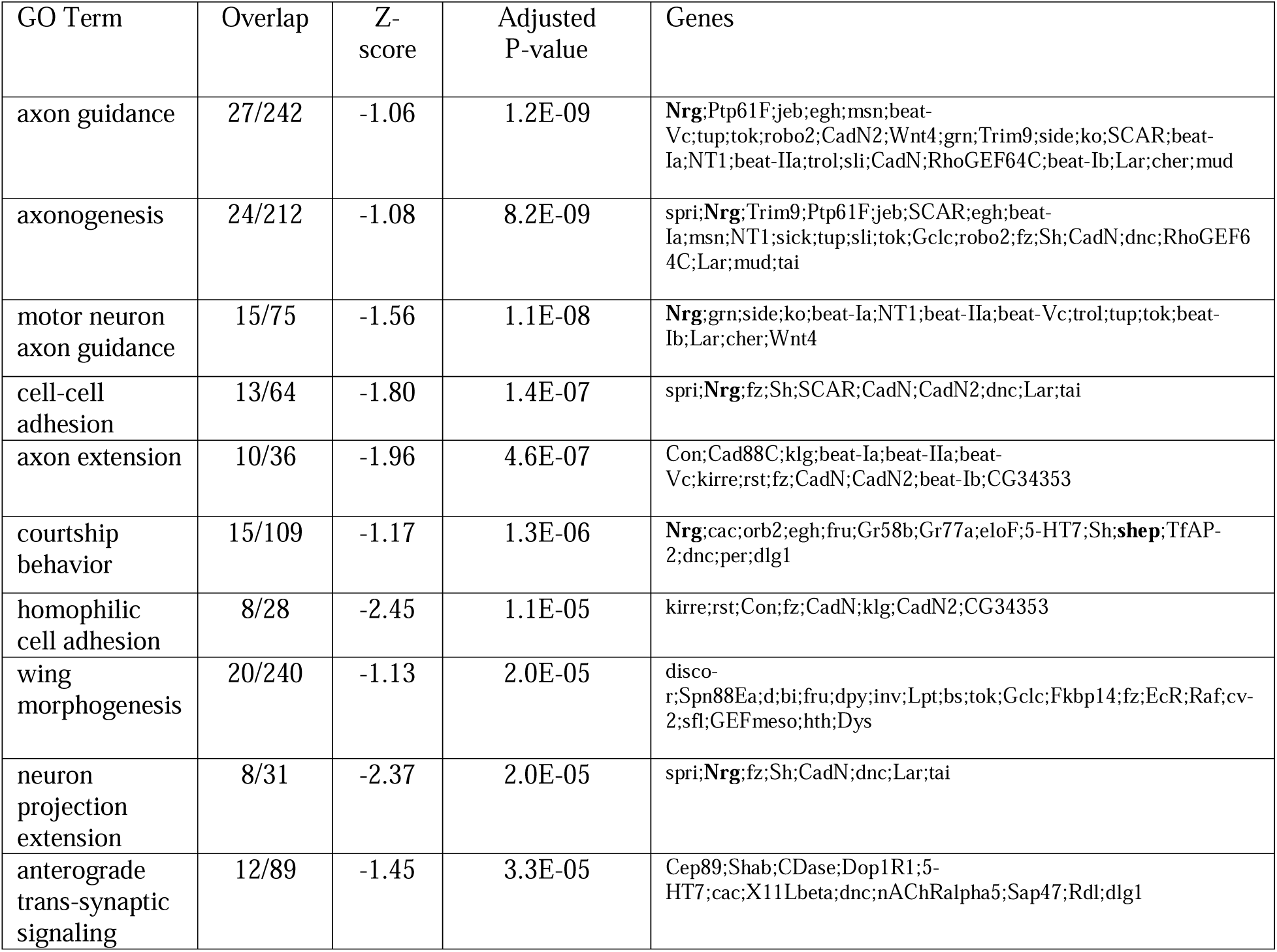
GO enrichment analysis using biological function highlights genes involved in neuronal development and behavior. Included are the top 10 GO terms with their representation in the data set and adjusted p-values after correcting for multiple testing.

We next compared our list of outliers to those outliers identified in Coughlan et al (2021). Coughlan et al (2021) conducted a GO enrichment analysis, with a different focus, that returned similar terms including behavior, mushroom body formation (neurogenesis), and olfaction as enriched terms (Fig 4 of Coughlan et al 2021). When comparing the list of outliers highlighted in that study (Supplemental Table 2) to our list of outliers we found five genes in common. These included *shep*, *Rdl*, *rad*, *RunxA*, and *Dop2R*. *Nrg* was in the top 5% of outliers from the Coughlan et al. (2021) study. In our final comparison we compared our list of outliers to genes that showed significantly different transcription in African and non-African female brains after exposure to both types of males (Bailey et al. 2011). In total there were 36 genes shared between these lists (Supplemental Table 3). The only gene that overlapped in all three studies was *shep*.

Given the patterns from GO enrichment analyses and comparisons with other data sets, we focused on *shep* and *Nrg* for functional tests. Previous behavioral analysis of homozygous viable mutant strains for these loci found evidence that these genes were involved female behavior (Carhan et al., 2005; Chen et al., 2014). Our goal was to use natural variation in *shep* and *Nrg* alleles to test for a role specifically in African female mate preference behavior.

### Population genetics and differentiation of shep and Nrg

Using previously published genomic datasets that sampled a large number of strains, we looked at patterns of nucleotide diversity (π) within two populations, patterns of differentiation (Fst) between these populations, and structure for *Nrg* and *shep*. π and Fst were summarized for 1 kb windows and compared to genome wide averages found in Lack et al (2016). For both *shep* and *Nrg* there was variability across the gene region for π and Fst (Table 2). While we would not necessarily expect the entire locus to be differentiated, *Nrg* had an average Fst that was greater than the genome wide average. The position of this locus on the X-chromosome could contribute to this pattern (reviewed in Meisel and Connallon 2013). Within both *shep* and *Nrg* there were windows with elevated Fst compared to the genome average. This differentiation was also captured by looking at the clustering of strains/genotypes in principal component space (Figure 2). The strains largely clustered by population for both loci. For *shep* there was also structuring within the ZI population, likely due to the presence of the segregating In(3L)P inversion in this region of the genome (Pool et al 2012; Corbett-Detig and Hartl 2012).

**Table 2.** The summary of population genetic statistics between the Raleigh (RAL) population, representing non-African populations, and Zambia (ZI), representing African populations, for *shep and Nrg*. Average values are provided for reference and come from Lack et al. (2016). We have provided averages for both the X and 3L chromosomes given their different effective population sizes. The mean across windows is provided with the (min, max) in parentheses.

| Gene | RAL pi | ZI pi | Fst |
| --- | --- | --- | --- |
| *shep*<br>(3L chromosome) | 0.0078<br>(0.0011,0.0202) | 0.0115<br>(0.0030,0.0253) | 0.1575<br>(0.0306, 0.4478) |
| *Nrg*<br>(X chromosome) | 0.0047<br>(0.0002,0.0140) | 0.0104<br>(0.0037,0.0262) | 0.2728<br>(0.0911,0.5788) |
| Averages | RAL average pi | ZI average pi | Average Fst |
| 3L Chromosome | 0.0065 | 0.0086 | 0.1811 |
| X Chromosome | 0.0041 | 0.0089 | 0.2708 |

### Shep contributes to female mate preference in an African strain

We used a set of crosses similar to quantitative complementation tests to determine if *alan shepard* (*shep*) contributes to female mate preference. We were able to validate that *shep* contributes to female mate preference by specifically demonstrating that females that carry only the African *shep* allele (African hemizygotes: A/N-*shep^-^*) prefer African males to non-African males (Figure 3A). When these data were analyzed using a binomial regression that included an interaction, the model did not contain any significant coefficients (Table 3). This is not unexpected given the specifics of our data and the phenotypes of F1s from our crosses where we did not *a* priori expect a hemizygote or strain effect. (See Methods and Supplemental Information for a full discussion). Given the clear trend for the African hemizygote female preference we used a model that directly estimated genotypes and was chosen specifically to reflect our expected phenotypes (see Methods and Supplemental Information). This analysis indicated that the African hemizygote females (A/N-*shep^-^*) were the only genotype that had a significant mating preference (Table 4). All other genotypes had 95% confidence intervals that contained zero indicating that they were consistent with random mating. We compared preference of the African hemizygote (A/N-*shep^-^*) to the preference of African heterozygote (A/N) using a Wald’s-test. This test indicated a significant difference in preference between these two genotypes (x^2^ = 13.0, *P* = 0.0015). Both genotypes are primarily heterozygous across the entire genome for non-African and African alleles since they are hybrids. The main difference between these genotypes is at the *shep* locus on chromosome 3, but there is the possibility, while unlikely, that other loci could contribute to this pattern (see Discussion)

**Figure 3.**
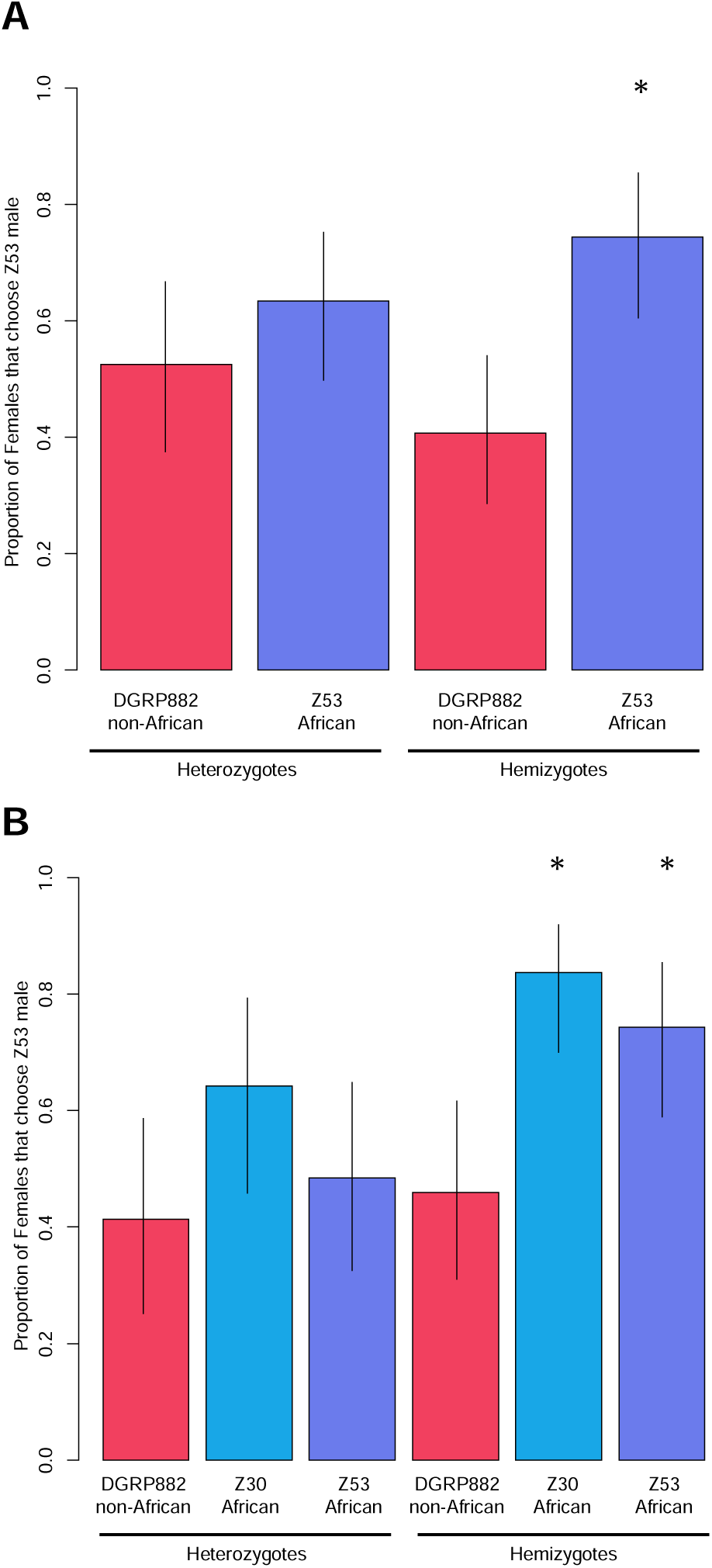
A) *alan shepard* (*shep*) and B) *Neuroglian* (*Nrg*) contribute to African female mate preference. Female flies that are hemizygous and carry only the African allele of *shep* and *Nrg* show increased preference for African males compared to heterozygote genotypes. This increase is in the direction of the wild-type African females. * represents a significant difference of the hemizygote genotype from the corresponding heterozygote genotype using a Wald’s test. Confidence intervals for each genotype are Wilson rank intervals estimated from the data.

**Table 3.** The analysis of female mating behavior using binomial regression containing interaction terms for both *shep* and *Nrg.* The African strain used in the cross is given next to the name of each locus. Coefficients represents the log-odds ratio for the specific effect/term. A discussion of the difference in these models and models without interaction terms is found in the Supplemental Information.

| ***shep* – Z53** |  |  |
| --- | --- | --- |
| Effect | Coefficient | *P*-value |
| $\mu$ - baseline | 0.1001 | 0.752 |
| $\beta_1$ – deficiency | -0.4748 | 0.259 |
| $\beta_2$ – strain | 0.4520 | 0.291 |
| $\beta_3$ - interaction | 0.9905 | 0.109 |
| ***Nrg* – Z53** |  |  |
| Effect | $\beta$ coefficient | *P*-value |
| $\mu$ - baseline | -0.3483 | 0.356 |
| $\beta_1$ – deficiency | 0.1858 | 0.711 |
| $\beta_2$ – strain | 0.2877 | 0.575 |
| $\beta_3$ - interaction | 0.9395 | 0.187 |
| ***Nrg* – Z30** |  |  |
| Effect | $\beta$ coefficient | *P*-value |
| $\mu$ - baseline | -0.3483 | 0.356 |
| $\beta_1$ – deficiency | 0.185 | 0.711 |
| $\beta_2$ – strain | 0.936 | 0.086 |
| $\beta_3$ - interaction | 0.864 | 0.255 |

**Table 4.** The behavioral differences in female mate preference for both *shep* and *Nrg* using models that compare each genotype to the random mating expectation. The coefficients represent log odds ratios, estimated from logistic regression, describing the increase or decrease in a genotype’s propensity to mate with African Z53 males. For this analysis we estimated 95% confidence intervals, and those that do not contain 0 are in bold.

| *shep* |  |  |
| --- | --- | --- |
| Strain-Genotype | Lower 95% CI | Upper 95% CI |
| DGRP882 heterozygote | -0.5222 | 0.7289 |
| DGRP882 hemizygote | -0.9300 | 0.1623 |
| Z53 heterozygote | -0.0018 | 1.1347 |
| Z53 hemizygote | **0.4141** | **1.7988** |
| *Nrg* |  |  |
| Strain-Genotype | Lower 95% CI | Upper 95% CI |
| DGRP882 heterozygote | -1.1117 | 0.3833 |
| DGRP882 hemizygote | -0.8204 | 0.4840 |
| Z30 heterozygote | -0.1662 | 1.3996 |
| Z30 hemizygote | **0.8901** | **2.5349** |
| Z53 heterozygote | -0.7523 | 0.6263 |
| Z53 hemizygote | **0.3800** | **1.8343** |

### Nrg contributes female mate preference in African strains

We used the same design to validate the role of *Nrg* alleles on female mate preference in African strains. Our design and logic were identical for what we used to analyze *shep*. For this gene we included more genotypes to determine if the visible marker on the FM7 balancer chromosome, *Bar*, which affects eye phenotypes was in turn affecting mating behavior. We also tested whether different African *Nrg* alleles might show different effects on female mate preference.

To test for the effects of the *Bar* allele specifically on female behavior we tested the behavior of another non-African strain, Canton S, and compared it to the DGRP882 strain. We used Canton S because we had a clear expectation: that both female genotypes from a cross between Canton S and the *Nrg^-^* strain should have the same behavior as the DGRP882 genotypes, and they should mate randomly and not show preference for either male genotype. We in fact observed that these genotypes mated indiscriminately, and we did not detect any significant difference in behavior comparing DGRP882 genotypes and Canton S genotypes (Supplemental Figure 3; Supplemental Table 5).

We then tested *Nrg* alleles from three African strains for effects on female mate preference. The first strain CH11 provided inconclusive results because females from both the heterozygote and hemizygote genotypes showed strong preference for Z53 males and had confidence intervals that were greater than 0 (Supplemental Figure 3; Supplemental Table 5). The dominant female preference from this strain precluded our ability to test the effect of the CH11 *Nrg* allele specifically.

The remaining two African strains had a pattern consistent with *Nrg* contributing to female mate preference (Figure 3B; Table 4). While binomial interaction models did not indicate a significant interaction affect (Table 3) our model that estimated the African hemizygote genotype suggested increased preference for the African hemizygotes (A/N-*Nrg*^-^) exclusively, as these genotypes were the only ones with 95% confidence intervals that did not contain 0 (Table 4). Additionally, a Wald’s test indicated a significant difference between African hemizygotes (A/N-*Nrg*^-^) and African heterozygotes (A/N) for both African strains that were tested 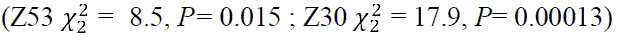. Similar to *shep,* there could be additional differences between the X chromosomes inherited by these genotypes (see Discussion), but the main difference is the *Nrg* alleles.

## Discussion

Identifying genes that contribute to female mate preference is an important first step in understanding how this important behavior evolves and contributes to the evolution of reproductive isolation and speciation. In this study, we demonstrate that *shep* and *Nrg* play a role in female mate preference in some African strains. Our current study is one of very few studies that has identified loci with direct genetic evidence for female mate preference (Chowdhury et al., 2020) and importantly in a system where premating isolation is the strongest and most relevant barrier to reproduction. These African populations of *D. melanogaster* have been a historically important system in the study of speciation (Wu et al., 1995; Coyne et al., 1999; Greenberg et al., 2003), and we have now identified, in part, genes that could contribute to premating isolation in this system. Since the genetic basis of behavior is complex, and we observed variation in behavior among strains (see below), it will be important to test the roles of *shep* and *Nrg* in more strains from different locations within Africa. Regardless, these remain promising candidates that we can leverage to understand the evolution of behavior. For example, when this *D. melanogaster* system was initially described, mapping studies were only able to identify large chromosomal regions on chromosome 3, the largest contributor to premating isolation, and the X chromosome (Hollocher et al., 1997; Takahashi and Ting, 2004).

Combining population genetics with complementation tests, we identified a gene on chromosome 3, *shep* is on 3L, and the X chromosome, *Nrg*, that contribute to female preference behavior. It is important to note that there could be additional differences on chromosome 3 and the X chromosome between the African hemizygote and African heterozygote because the chromones they inherited from the mutant/balancer strain are non-recombining (Supplemental Figure 1). While this lack of recombination could lead to the accumulation of additional differences between these chromosomes, having any mutations result in the specific phenotype for increased female preference when combined with an African genome would be unlikely. One reason is that even if these chromosomes were different, they are still more similar in terms of behavioral alleles (both are non-African) to each other than either are to African chromosomes. If there were differences, they would either need to be null mutations in a behavioral gene, which would create a hemizygote similar to the gene we are testing, or have some epistatic interaction with the African genome. Considering both of these factors together it is more likely that *shep* and *Nrg* alleles are responsible for our signal for female preference.

Getting gene scale resolution will allow us to further understand how female mate preference has evolved. Since both *Nrg* and *shep* are essential for neurodevelopment and female mating behavior (Carhan et al., 2005; Chen et al., 2014), the identification of their role in female preference in African strains can potentially hint at sensory pathways that are important for mating decisions and partial reproductive isolation. Female preference can evolve when structures/genes that are involved in sensory perception evolve, when brain centers important for learning/decision making evolve, or when both evolve in tandem (Stevens, 2013; Schaefer and Ruxton, 2015; Munson et al., 2020). While we focused on *Nrg* and *shep* because they were identified as outliers or genes of interest in several independent analyses, future work could also follow up on additional genes identified in our analyses that are involved in neurodevelopment. For example, *Shaker (Sh)* was represented in several GO terms and has known roles in neurotransmission and mating behavior (Pimentel et al 2016). Many studies have documented the role that female preference plays in reproductive isolation (Laturney and Moehring 2012), and further demonstrating how sexual selection can shape evolution of female preference and the genetic architecture of female preference (Xu and Shaw 2021). Both *shep* and *Nrg* regulate neurological development, specifically neural remodeling during metamorphosis, contributing heavily to the formation of higher learning centers, specifically those responsible for processing olfactory signals (Carhan et al., 2005; Chen et al., 2014). Our gene ontology results highlight a role for genes involved in neurogenesis and behavior that complements other gene ontology analyses that suggest outlier genes are involved in sensory perception and neurological development.

Previous work in African populations of *D. melanogaster* has suggested a large role for olfaction in female mate preference (Grillet et al., 2012; Moran, 2006; Jin et al., 2022) and it is possible that *Nrg* and *shep* may contribute to these observed patterns. *Nrg* controls, in part, the development of the mushroom body (Goossens et al 2011) which is a structure in *Drosophila* that processes olfactory information (Li et al 2020). It is possible that differences in *Nrg* across populations could result in differences in the mushroom body, either at the gross anatomical level, or at the circuit level and how it receives input from olfactory receptors (Akalal et al. 2006). In closely related species of *Drosophila* differences in olfactory preference can be explained by the connections made between olfactory receptors and the mushroom body (Ellis et al. 2023). *shep* is an RNA/DNA binding gene that regulates alternative splicing (Chen et al. 2018; Olesnicky et al. 2018). Given the evidence for changes in *shep* expression in African and non-African strains under different mating conditions (Bailey et al. 2011), shep might regulate targets in specific neurons that contribute to mating behaviors. It has been demonstrated that *shep* has specific targets during neuronal remodeling (Chen et al 2018) and might interact with this or other targets in the adult brain (Olesnicky 2018). Polymorphisms in the coding regions of either gene could alter interactions with other genes/targets during neuronal development.

Changes in regulatory regions and/or introns could change expression patterns. More work is needed to establish functional differences between African and non-African alleles. Specifically, whether functional changes in the neural networks of females are correlated with the olfactory cues used in mate preference. Regardless, while we observed large effects of these single loci, the genetic basis of female preference is likely complex, with additional loci contributing to isolation in these strains.

One interesting observation that could suggest that more than these two loci contribute to female preference is the variation in dominance that we observed for female mating preference in hybrids between non-African and African strains. In our experiment, we used three African strains that show strong preference for African males and never choose non-African males in choice tests. We observed that female mate preference was recessive in F1 hybrids for two of the strains and dominant in F1 hybrids the third strain when crossed to the same non-African strain (see Results). The dominance in the CH11 strain precluded us from testing the effects of *Nrg* on female preference for that strain. When assessing how important *shep* and *Nrg* are in reproductive isolation across strains different mutations (i.e., CRISPR knockouts) can be used to circumvent this issue. Nevertheless, testing both the effects of *shep* and *Nrg* and the dominance of female behavior in more strains will be informative in and of itself and could highlight how general this phenomenon is. This will be important for two reasons. First, if more strains show effects of *shep* and *Nrg* on female preference this could suggest these genes are critical for reproductive isolation in the system (see below). However, it should be cautioned that at early stages of divergence we do not necessarily expect fixed differences between populations (Cutter, 2012; Castillo and Barbash, 2017). This contrasts with reproductive isolation between species where we would assume genetic differences are fixed (Laturney and Moehring 2012). Second, in terms of dominance, looking for general patterns will be important because the dominance of reproductive barriers can have strong impacts on the outcome of speciation (Thompson et al., 2021) and might indicate how particular mating barriers evolve.

Differences in dominance for female mate preference could reflect differences in the selective environment caused by local mating dynamics. This scenario would be analogous to parallel vs non-parallel evolution of ecological traits in different environments (Oke et al., 2017; Bolnick et al., 2018). If different populations experience different selective pressures, that is there is variation in male courtship behavior and female preference across populations, then different alleles of *shep* and *Nrg* could segregate in these populations. African lineages are quite diverse in terms of courtship behaviors (Colegrave et al 2000; Yukilevich and True 2008; Jin et al. 2022) and while strong premating isolation is found in many populations it is likely that female preferences and associated alleles could be segregating within Africa (Yukilevich and True, 2008; Coughlan et al 2021). Another possibility, that is not mutually exclusive, is that different genes contribute to female mate preference in different populations. This would generate a pattern where genes show effects on behavior for some hybrid genotypes, but not others. Given the limited data on the genetic basis of female preference it is difficult to infer the likelihood of this process, but polymorphic incompatibilities are common in other systems (Cutter, 2012; Castillo and Barbash, 2017).

One example in *Drosophila* suggests that the genetic architecture of premating isolation can be variable. When different strains of *D. simulans* were crossed and tested for mate rejection of *D. melanogaster,* different numbers, locations, and dominance of quantitative trait loci were identified that were strain specific (Uenoyama and Inoue, 1995; Carracedo et al., 1998a; Carracedo et al., 1998b; Carracedo et al., 2000). The variation in dominance and the number of loci contributing to mate preference in *D. simulans* might be consistent with this barrier evolving after the speciation event that separated these lineages (Laturney and Moehring, 2012). Moving forward it will be important to determine how common it is for variable loci to contribute to premating isolation or uniform genetic architecture for reproductive isolation between populations. Either way, the difference in dominance suggests there may be standing variation for female mate preference that selection can act on, which is known to facilitate the evolution of premating isolation through sexual selection (Mendelson et al., 2014; Castillo and Delph, 2016).

Overall, our results provide evidence for two neurodevelopmental genes, *Nrg* and *shep,* contributing to female mate preference in African strains, suggesting that these genes could contribute to mating preferences between populations of *D. melanogaster*. Our results connect population genetics from a behavioral cline, to allelic diversity in African and non-African populations, to functional validation of the effect of these genes on female African preference behavior. The differences in alleles for both loci in non-African vs African populations should be further explored and the testing of more strains are needed to determine whether these genes could contribute to reproductive isolation. Understanding the functional consequences of different alleles on neural circuits and sensory perception for specific alleles could provide insight into how female preference evolved in this system, and provide a framework for exploring these types of data in other systems. Ultimately, this could lead to understand common patterns for the evolution of female mate preference and the genetic and neural level.

