## Supplemental Information for "The neurodevelopmental genes *alan shepard* and *Neuroglian* contribute to female mate preference in African *Drosophila melanogaster*"

**Suppleme­­­­­­­­ntal Information**

**Testing for impacts of food-dye on female choice experiments**

In a preliminary experiment, using strains Z53 and DGRP882, we conducted 30 mate choice trials all within strain, and there was no effect of blue food coloring on female preference. We analyzed data using an exact binomial test with the null hypothesis that the proportion was equal to 0.5. Z53 females chose blue-dyed males 14 out of 30 times (*P*= 0.8555), and DGRP882 females chose blue-dyed males 17 out of 30 times (*P*=0.5847). We then tested whether coloring affected mating preferences between strains Z53 and DGRP882 and found no significant effect. In crosses with DGRP882 females 15 of the crosses contained DGRP882 dyed blue paired with non-dyed Z53, and 15 crosses contained Z53 dyed blue with non-dyed DGRP882. In these crosses blue dyed males (ignoring genotype) were chosen 15 out of 30 times (*P*=1.000), and DGRP882 males were chosen 16 out of 30 times (*P*=0.8555). In crosses with Z53 females DGRP882 males were never chosen (0 out of 30) regardless if they were blue or non-dyed, and Z53 males were always chosen (30 out of 30). Because there were 15 trials with Z53 blue, blue was chosen 15 out of 30 times.

**Statistical Analysis**

In a standard complementation test, the starting point is two parental strains that have divergent phenotypes for a continuous trait, and are often the minimum and maximum trait values in a population/species (reviewed in McKay 2010). To analyze the differences in these genotypes a linear model of the following structure is used (Equation 1). The goal is to isolate the effects of the gene of interest on the phenotype while taking into account the difference in parental strains and any effects the balancer chromosome might have on the phenotype.

$y= \mu+\beta_{1}x_{1}+\beta_{2}x_{2}{+\beta}_{3}x_{1}x_{2}$	Equation 1

The design matrix is structured such that x1=0 when the genotype does not carry the deficiency of the gene/locus of interest (and therefore carries the balancer chromosome) and x1=1 when the genotype does carry the deficiency (and lacks the balancer chromosome). For x2, x2=0 when the genotype comes from the P1 strain and x2=1 when the genotype comes from the P2 strain with the divergent trait value. For the model: μ is the intercept, β_1_ is the effect of carrying a deficiency for the locus of interest, β_2_ is the strain effect, and β_3_ is the interaction effect and the change in phenotype when the genotype is both from the P2 strain and carries the deficiency. How these coefficients correspond to the expected phenotypes from a quantitative complementation tests are depicted in Supplementary Figure 2.

The genotype mean trait values from the model would be:

Genotype 1: P1 heterozygote (baseline) = μ

Genotype 2: P1 hemizygote, deficiency effect but no strain effect = μ + β_1_

Genotype 3: P2 heterozygote, strain effect but no deficiency effect = μ + β_2_

Genotype 4: P2 hemizygote, strain and deficiency effect = μ + β_1_ + β_2_+ β_3_

The genotypes that carry the deficiency are hemizygous. That is, they only carry one allele, the parental wild-type allele, for the gene of interest. If this gene contributes to the phenotype the expectation is that the hemizygotes will have a phenotype that resembles the respective parental strains (who would presumably be homozygous for this allele).

The interpretation of the interaction term is important in these complementation tests as often the main effects (β_1_ and β_2_) have opposite signs (Supplemental Figure 2). While this is not a requirement, it is often a consequence of the structure of the experimental crosses and statistical model. The Genotype equations above can be used to understand this logic. If the P1 parent produces heterozygotes and hemizygotes that have smaller trait values then β_1_ will be negative because Genotype 2 will have a smaller trait value compared to Genotype 1. The P2 strain would have a higher trait value as a heterozygote (Genotype 3) and β_2_ would be positive. Therefore, to conclude there is an effect of the gene of interest, the interaction term (β_3_) would need to be significant and Genotype 4 would have a trait value greater than Genotype 3. Significant main effects and no interaction effect would indicate only a strain effect. In the absence of main affects (β_1_=0, β_2_=0), then only β_3_ would be informative. However, this would not be expected under the typical complementation test where hybrids are intermediate in phenotype and hemizygote genotypes (Genotype 2 and Genotype 4) would have trait values shifted in opposite directions from F1 heterozygotes (Supplemental Figure 2). Additionally, you would always expect a strain effect because you chose the parental strains based on their trait difference before conducting crosses.

*Binomial Regression – Interaction Model and Alternatives*

In our preference experiments the data can be represented as binary choices. An individual female replicate chooses either the DGRP882 or the Z53 male, which can be represented as 0 and 1. Our statistical philosophy is to model the data in its natural form (Warton et al. 2016) and would lead us to binomial regression. One benefit of the generalized linear models is that the same model structure can be used as Equation 1, except now the scale is in log-odds due to the logit transformation of the data. The logit link function is the most common link function for binomial regression (Equation 2).

$logit(p)= \mu+\beta_{1}x_{1}+\beta_{2}x_{2}{+\beta}_{3}x_{1}x_{2}$	Equation 2

The interpretation of the model is the same as presented for the linear model above. While this approach conveniently matches the rich literature of complementation tests, there are some barriers to using this model for our experiment. For a binomial variable representing choice, the maximum would be a genotype never choosing one male (0 %) and the opposite genotype choosing the other male exclusively (100%). In our data we instead have female genotypes that mate at random and choose one male at 50% and choosy females that choose that same male 100% of the time. As we describe in the methods of the main text this creates a difficulty because our random mating parental strain has the same phenotype as any mutant/balancer strain used. As a result, F1s produced from crosses between these strains are expected to have a similar phenotype (Supplementary Figure 2). This results in a scenario where we are unlikely to see shifts in hemizygous genotypes in opposite directions which is a requirement for “failure to complement” in a standard quantitative complementation test.

There are two examples from the literature worth examining and comparing with our data and analyses since both use quantitative complementation tests but have different phenotype expectations. The study of Chowdhury et al., (2020) uses parental strains (*D. simulans* and *D. melanogaster*) that have completely divergent behavior phenotypes in their assays. In the crosses of these parentals with the mutant/balancer strain they have expectations that match panels A-C in Supplementary Figure 2. They use the binomial regression with the interaction term and observe significant interaction when the hemizygous genotypes had shifts in phenotypes in the opposite direction from one another (ie toward respective parental genotypes). It is worth noting that they report the significance of the interaction but not the direction or re-calculate the probabilities to demonstrate the sign change for each genotype. This is typically not sufficient to understand the sign of the interaction effect without recalculating the odds ratios for the comparisons of interest (Ai and Norton 2003; Chen 2003). The interpretation of interaction effects in logistic regression cannot be made based on this single coefficient because the interaction is conditional on other effects (Ai and Norton 2003).

In the second study that uses quantitative complementation tests and binomial regression, Comeault et al. (2017) utilize parental strains that are divergent (*D. melanogaster* and *D. orena*), but the behavior of the mutant/balancer strain is identical to the wild-type *D. melanogaster* parental strains (since they are both *D. melanogaster*). In this study they are testing the effects of olfactory receptors on the use/preference for waterberry in *D. orena*. This fruit does not elicit a response in any *D. melanogaster* strains. This creates a scenario akin to panel D-F in Supplemental Figure 2 and similar to our study. Because of the shared phenotype for *D. melanogaster* parental and mutant/balancer strains you do not expect an overall hemizygosity effect. In fact, Comeault et al (2017) only report crosses for heterozygotes and hemizygotes that use the *D. orena* parent and only make inference based on comparing these two genotypes.

**Supplemental Tables**

**Supplemental Table 1.** Genes that had a SNP within the gene that was an outlier at the 1% threshold in our cline analysis. This list was used for GO enrichment analysis and comparisons with other datasets.

| Gene name | | | | | | | | |
| --- | --- | --- | --- | --- | --- | --- | --- | --- |
| *2mit* | *CG12096* | *CG32686* | *CG7029* | *Dhc98D* | *hth* | *P58IPK* | *SIFaR* | *Vps4* |
| *4EHP* | *CG12125* | *CG3277* | *CG7058* | *dikar* | *iab-8* | *PAPLA1* | *sima* | *Vrp1* |
| *5-HT7* | *CG1273* | *CG32809* | *CG7208* | *disco-r* | *iav* | *pav* | *SkpD* | *Vsx1* |
| *Ac13E* | *CG13085* | *CG32982* | *CG7376* | *dlg1* | *Ilp6* | *Pburs* | *slam* | *wake* |
| *ACXD* | *CG13252* | *CG33060* | *CG7432* | *Dnah3* | *inv* | *pcx* | *sli* | *wat* |
| *Afti* | *CG13615* | *CG33090* | *CG7708* | *dnc* | *Jarid2* | *Pde9* | *SmydA-8* | *wb* |
| *aPKC* | *CG13842* | *CG33181* | *CG7804* | *Dop1R1* | *jeb* | *per* | *sNPF* | *Wdr62* |
| *app* | *CG14120* | *CG33298* | *CG7886* | *Dop2R* | *kar* | *Pex5* | *sofe* | *whd* |
| *Arp10* | *CG14204* | *CG33958* | *CG7888* | *dpr1* | *kirre* | *pico* | *Spn88Ea* | *Wnt4* |
| *Arp5* | *CG14218* | *CG34260* | *CG7985* | *dpr8* | *kkv* | *Piezo* | *SppL* | *wry* |
| *asp* | *CG14219* | *CG34347* | *CG8562* | *dpy* | *klg* | *PIG-S* | *spri* | *X11Lbeta* |
| *Atf3* | *CG14621* | *CG34353* | *CG8563* | *dsb* | *Klp67A* | *PIG-U* | *SRm160* | *boss* |
| *Atg2* | *CG14778* | *CG34362* | *CG8907* | *Dys* | *ko* | *Plod* | *ssp3* |  |
| *ATP8B* | *CG14913* | *CG34370* | *CG9265* | *e(y)1* | *kz* | *poly* | *Strn-Mlck* |  |
| *b6* | *CG14915* | *CG34383* | *CG9518* | *Eaat2* | *l(2)35Df* | *prd* | *Sur* |  |
| *bab1* | *CG15312* | *CG34417* | *CG9815* | *EcR* | *Lap1* | *Proc* | *Syn2* |  |
| *Baldspot* | *CG15431* | *CG3797* | *CG9850* | *Efr* | *Lar* | *Proc-R* | *Syp* |  |
| *beat-Ia* | *CG15465* | *CG4004* | *CG9970* | *egh* | *Liprin-gamma* | *Prosap* | *Syt4* |  |
| *beat-Ib* | *CG15570* | *CG42331* | *chas* | *egl* | *Lkr* | *ps* | *Sytbeta* |  |
| *beat-IIa* | *CG15618* | *CG42594* | *cher* | *eIF-4a* | *Lmpt* | *Ptp61F* | *T48* |  |
| *beat-Vc* | *CG15625* | *CG42660* | *ChLD3* | *Eip63E* | *Lpt* | *pxb* | *tai* |  |
| *Best2* | *CG1635* | *CG42661* | *cindr* | *Elk* | *LRP1* | *pyr* | *Tao* |  |
| *bi* | *CG16771* | *CG42666* | *Cirl* | *eloF* | *luna* | *rad* | *TBC1D5* |  |
| *boss* | *CG16791* | *CG42748* | *Clamp* | *et* | *mamo* | *Raf* | *TfAP-2* |  |
| *br* | *CG16833* | *CG42749* | *cmpy* | *Evi5* | *mask* | *Rdl* | *Tie* |  |
| *bru-3* | *CG17097* | *CG43175* | *Con* | *fest* | *Mct1* | *rg* | *timeout* |  |
| *bs* | *CG17129* | *CG43394* | *CR32773* | *Fili* | *MED10* | *Rgk1* | *tinc* |  |
| *btsz* | *CG17186* | *CG43427* | *CR33013* | *Fkbp14* | *MED28* | *rgn* | *Tl* |  |
| *Ca-alpha1T* | *CG17841* | *CG43693* | *CR43637* | *Flo2* | *mRpS5* | *rhea* | *Tm1* |  |
| *cac* | *CG1791* | *CG43736* | *CR43751* | *fru* | *ms(2)35Ci* | *RhoGAP19D* | *TM9SF4* |  |
| *Cad88C* | *CG18031* | *CG43783* | *CR44172* | *fs(1)K10* | *msi* | *RhoGEF64C* | *tn* |  |
| *CadN* | *CG2121* | *CG44085* | *CR44347* | *Fur1* | *msn* | *Rip11* | *Tnks* |  |
| *CadN2* | *CG2528* | *CG44422* | *CR44350* | *fz* | *MsR1* | *robo2* | *tok* |  |
| *Calx* | *CG2772* | *CG45186* | *CR44563* | *fzr2* | *mthl11* | *rst* | *Tpr2* |  |
| *car* | *CG2970* | *CG4607* | *CR44592* | *Galphaf* | *mud* | *RunxA* | *Trim9* |  |
| *caz* | *CG30043* | *CG4751* | *CR44636* | *Gclc* | *Mur2B* | *Samuel* | *trol* |  |
| *CCHa1* | *CG3091* | *CG4928* | *CR44754* | *GEFmeso* | *mura* | *sano* | *trpl* |  |
| *CDase* | *CG3106* | *CG4972* | *CR44811* | *Gfrl* | *nAChRalpha5* | *Sap47* | *Tsp39D* |  |
| *CenG1A* | *CG31268* | *CG4991* | *CR44846* | *GluRIB* | *nahoda* | *Sas10* | *Tsp66A* |  |
| *Cep89* | *CG31300* | *CG5080* | *CR45109* | *GlyP* | *Ndae1* | *SCAR* | *Tsp66E* |  |
| *CG10011* | *CG31327* | *CG5160* | *CR45190* | *GNBP2* | *NK7.1* | *sev* | *tud* |  |
| *CG10163* | *CG31446* | *CG5196* | *CR45753* | *goe* | *Nlg2* | *sfl* | *tup* |  |
| *CG10514* | *CG31663* | *CG5278* | *CR45996* | *Gr58b* | *Nrg* | *Sfmbt* | *TwdlG* |  |
| *CG10543* | *CG31874* | *CG5382* | *CR46033* | *Gr58c* | *nrm* | *Sgs5* | *TyrRII* |  |
| *CG10839* | *CG3191* | *CG5565* | *CR46264* | *Gr77a* | *Nrx-IV* | *Sh* | *upd2* |  |
| *CG1124* | *CG32206* | *CG5604* | *cv-2* | *Grip84* | *NT1* | *Shab* | *ush* |  |
| *CG11317* | *CG32302* | *CG5955* | *Cyp301a1* | *grn* | *Obp19c* | *shakB* | *VGlut* |  |
| *CG11321* | *CG32365* | *CG6006* | *d* | *Gyc32E* | *Or2a* | *shep* | *Vha100-3* |  |
| *CG11873* | *CG32533* | *CG6540* | *Dat* | *HBS1* | *orb2* | *sick* | *Vinc* |  |
| *CG11905* | *CG32683* | *CG7025* | *Dhc16F* | *HDAC4* | *osp* | *side* | *Vps16A* |  |

**Supplemental Table 2**. Outliers highlighted in Coughlan at al (2021) Figure 4 that we compared to our list of outliers

| *orb22b* | *beat-Illc* | *CG5050* | *lig* | *ths* | *Dpr13* | *shep* |
| --- | --- | --- | --- | --- | --- | --- |
| *glut1* | *hng3* | *TSG101* | *Crtc* | *sif* | *vn* | *Or67c* |
| *Ank2* | *CKIIIalpha* | *Rbp6* | *Rdl* | *CG7084* | *Ir94* | *Pzi* |
| *OR85e* | *CG5921* | *Grip* | *DAAM* | *rg* | *Tbh* | *rad* |
| *RunxA* | *Dop2R* | *eag* |  |  |  |  |

**Supplemental Table 3.** The intersection of genes that were outliers in our study and in Bailey et al (2011) that analyzed changes in expression within female brains.

| *CG33090* | *Vinc* | *mask* | *RhoGAP19D* | *l(2)35Df* | *Prosap* | *Baldspot* |
| --- | --- | --- | --- | --- | --- | --- |
| *CG4751* | *Sfmbt* | *timeout* | *Dys* | *kz* | *Lar* |  |
| *osp* | *rhea* | *fz* | *pcx* | *Sas10* | *X11Lbeta* |  |
| *Tsp39D* | *SCAR* | *Lap1* | *asp* | *CG33298* | *sfl* |  |
| *egh* | *dlg1* | *cher* | *CG12125* | *mas* | *dpr8* |  |
| *CG33181* | *Nrx-IV* | *shep* | *msi* | *Tm1* | *CG15570* |  |

**Supplemental Table 6.** The mating preference for Canton S and CH11 genotypes when tested for the effect of *Nrg* on behavior. Canton S was used to test whether any effect of the *Bar* mutation. We expected no difference in behavior compared to DGRP882 if *Bar* did not affect preference and confirmed this. For CH11 we were unable to test the effect of *Nrg* since African female preference was dominant in these hybrids and we could not differentiate the heterozygotes from the hemizygote preference. For this analysis we estimated 95% confidence intervals, and those that do not contain 0 are in bold.

| ***Nrg*** | | |
| --- | --- | --- |
| Strain-Genotype | Lower 95% CI | Upper 95% CI |
| DGRP882 heterozygote | -1.111 | 0.383 |
| Canton S heterozygote | -0.637 | 0.989 |
| CH11 heterozygote | **0.518** | **2.745** |
| DGRP882 hemizygote | -0.820 | 0.484 |
| Canton S hemizygote | -0.523 | 0.957 |
| CH 11 hemizygote | **0.556** | **2.379** |

**Supplemental Figures**

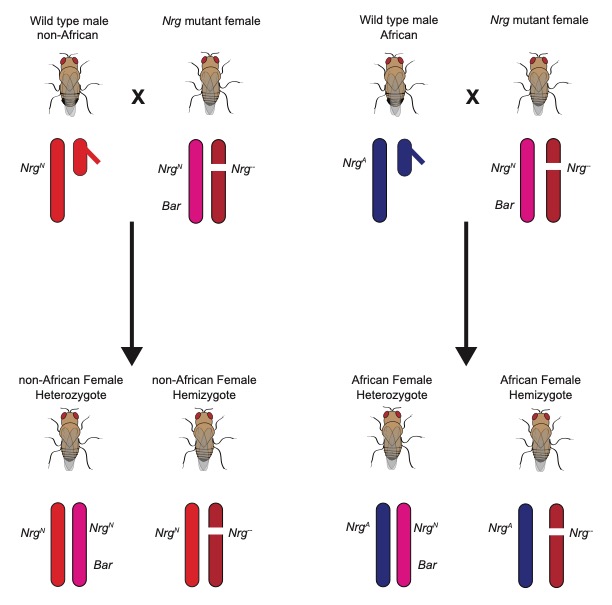

**Supplemental Figure 1.** A diagram depicting the crossing scheme used for quantitative complementation tests. This specific diagram illustrates the design for *Nrg*. The design for shep was identical, but *shep* is on the third chromosome whereas *Nrg* is on the X chromosome. *Nrg^N^* represents the non-African allele of *Nrg.* *Nrg^A^* represents the African allele and *Nrg^-^* represents the null allele. The distance/orientation of *Nrg* and *Bar* is not to scale, but just to illustrate the linkage on the FM7 balancer (pink) chromosome. Each chromosome is a different color, but the red and pink shades are more similar because they come from non-African origin.

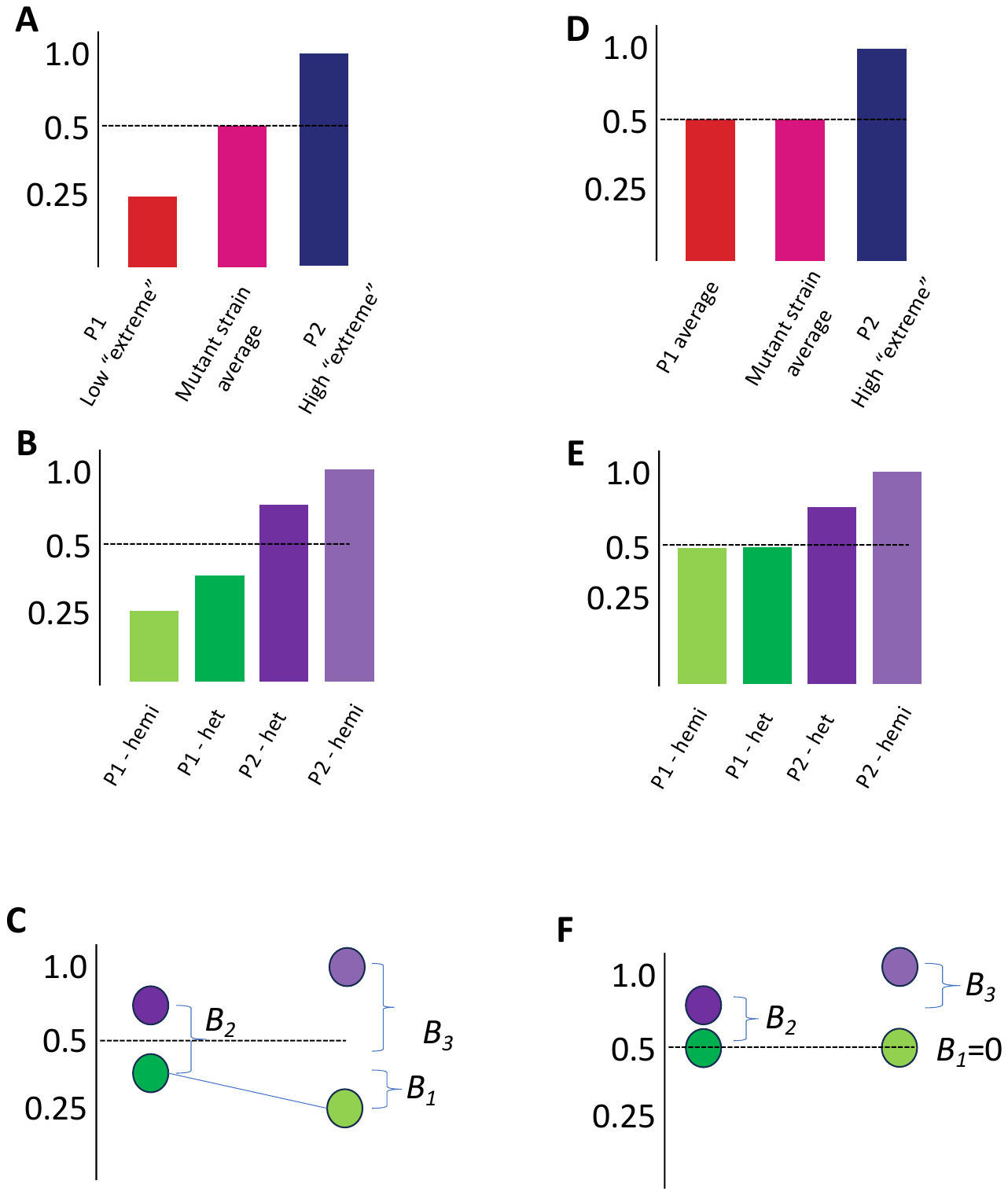

**Supplemental Figure 2.** A-C represent the expected phenotypes for parental strains and F1 progeny from a standard quantitative complementation test that uses binary choice assays. Panels D-F. Represent a scenario closer to what is observed in this study. Specifically, there cannot be divergent “extreme” parent values with and average mutant/balancer strain. Since all mutants are from outside of Africa then there is a lower limit on what we will observe. We will never observe female preference lower than 50%. The Y axis for all panels is the proportion of females that choose a specific male genotype. A) Parental strains have divergent phenotypic values and are crossed with a mutant/balancer strain that has an “average” phenotype with respect to the entire population/species. B) The F1 are represented as showing deviations from 50% due to both strain effects and hemizygosity effects. C) The same data are represented/reorganized to highlight the connection between specific genotypes and the regression coefficients (similar to reaction norms). The result is that the alleles fail to complement indicating the gene being tested contributes to the phenotype. If the P1 het is baseline (mu), then *B1* is the effect of the hemizygote, *B2* is the effect of the P2 strain, and *B3* is the interaction term F). S As a result, all coefficients have decreased in magnitude. This means that there is no hemizygote effect (B1=0). Now both B2 and B3 are smaller and potentially harder to estimate. Note that the difference between P2-F1 genotypes remains unchanged.

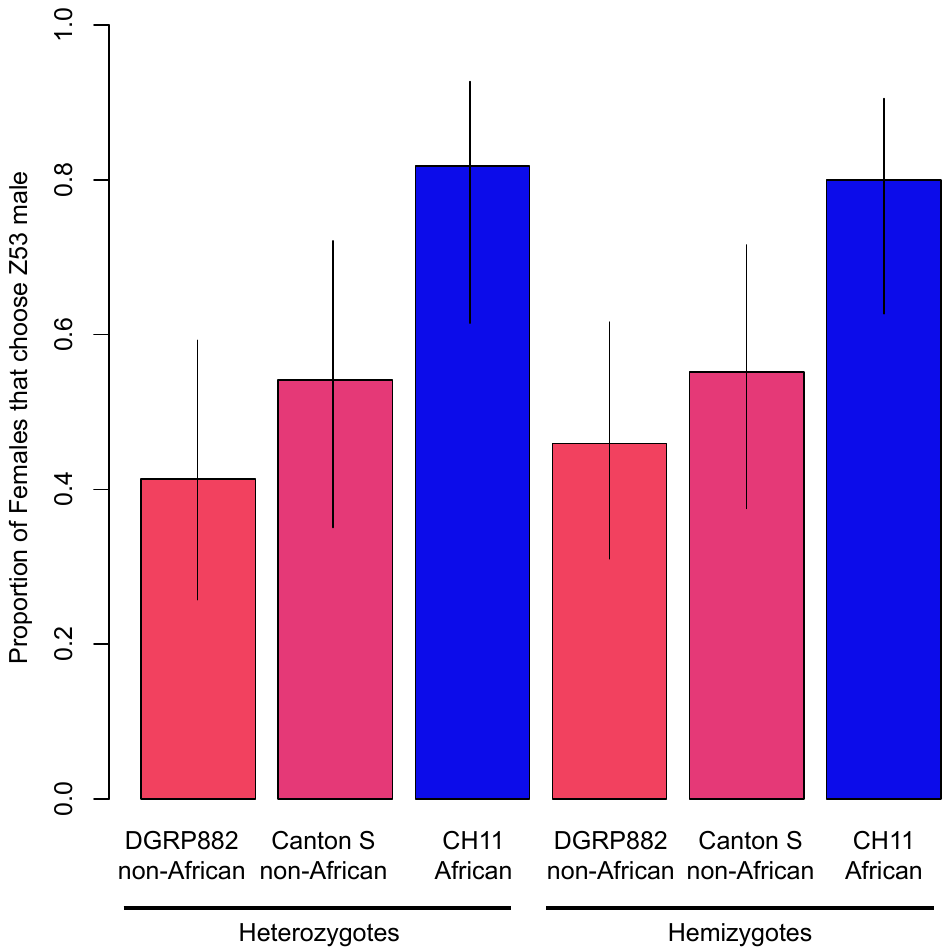

**Supplemental Figure 2**. The dominant visible marker, *Bar*, on FM7 does not affect female mate preference for either non-African or African strains. Canton S was included as a control because we expected it to behave similarly to the DGRP882 strain. The CH11 strain was included to test multiple African *Nrg* alleles. Because this strain exhibited a dominant female preference phenotype for both heterozygous and hemizygous females these genotypes were indistinguishable from each other. Confidence intervals are Wilson rank intervals estimated from the data.
